# Laudanosine restricts Ebola virus entry by targeting TPC2-dependent endolysosomal trafficking

**DOI:** 10.64898/2026.06.17.732874

**Authors:** Tom Seitz, Julian Koengeter, Susanne Klute, Franziska Kraft, Denise C. Clesle, Lily Tschampel, Nico Preising, Armando Alexei Rodríguez Alfonso, Sebastian Wiese, Ludger Ständker, Christoph Jung, Timo Jacob, Janet Köhler, Gilbert Weidinger, Nadine Biedenkopf, Konstantin M.J. Sparrer, Frank Kirchhoff, Fabian Zech

## Abstract

Late endosome-dependent viruses, including filo- and arenaviruses, rely on host endolysosomal trafficking for productive infection. Here, we used a dual-colour Vesicular stomatitis virus (VSV) based pseudoparticle screen of CytoSorb-derived fractions to identify inhibitors of the Zaire Ebolavirus glycoprotein (GP)-mediated entry. Iterative chromatographic purification and mass spectrometry identified Laudanosine, a degradation product of the clinically used neuromuscular blocker Atracurium, as the antiviral compound. Laudanosine specifically inhibited entry mediated by Ebola, Marburg, Lymphocytic choriomeningitis and Lassa virus glycoproteins without affecting VSV-G-dependent entry. Importantly, Laudanosine inhibited authentic Ebola virus infection without detectable cytotoxicity in cell culture and embryonic zebrafish. Molecular dynamics simulations suggest stable association of Laudanosine with the allosteric inhibitory pocket of the lysosomal two-pore channel (TPC2). Consistently, Laudanosine impairs autophagic flux and disrupts endolysosomal trafficking. Together, our findings identify Laudanosine as a previously unrecognised inhibitor of TPC2-dependent entry of highly lethal viral pathogens.

## INTRODUCTION

Viral pathogens continue to pose major threats to global health because therapeutic options remain limited^1^. For many clinically relevant viruses, including highly lethal filoviruses and arenaviruses such as Ebola and Lassa viruses, broadly effective drugs and vaccines are missing^2^. Thus, innovative strategies to identify antiviral agents, including compounds targeting conserved host-dependent processes required for viral replication, are urgently needed.

Human body fluids and tissues represent a rich and largely unexplored source of naturally occurring bioactive molecules that may contribute to intrinsic antimicrobial defence^3,4^. We have previously exploited peptide libraries generated from human-derived materials to identify novel antiviral factors. This strategy enabled the discovery of endogenous peptidic inhibitors targeting HIV-1, cytomegalovirus (CMV), and SARS-CoV-2 infection, several of which show promising properties for further development as antiviral therapeutics^5–8^. Beyond peptide inhibitors, body-fluid-derived libraries can also reveal broader antiviral mechanisms. For example, phosphatidylserine-exposing extracellular vesicles present in human semen and saliva have been identified as an innate defence against viral pathogens using apoptotic mimicry for attachment and entry^9^.

Host-directed antivirals are attractive against viruses that depend on specific cellular pathways for infection. Late endosome-dependent viruses, including filoviruses and arenaviruses, require coordinated intracellular trafficking events to reach compartments permissive for membrane fusion and viral entry. Two-pore channels (TPCs), located in late endosomal and lysosomal membranes, play a central role in these processes^10,11^. Upon activation by nicotinic acid adenine dinucleotide phosphate (NAADP), TPCs regulate endolysosomal ion homeostasis and promote vesicular maturation and fusion events^12,13^. Ebola virus, Marburg virus, Lassa virus and related pathogens exploit TPC-dependent trafficking pathways during entry. Pharmacological inhibition of these channels traps incoming viral particles in non-productive compartments^14,15^. Thus, TPC-regulated endolysosomal trafficking represents a promising target for antiviral intervention, as its inhibition halts lysosomal fusion, blocks autophagic flux, and prevents the release of viral genomes into the cytoplasm.

Vesicular stomatitis virus (VSV)-based pseudoparticles (pp) lacking the endogenous glycoprotein (VSVΔG) provide a versatile platform to identify entry inhibitors against highly virulent viral pathogens under reduced biosafety conditions^16^. In this study, we established a high-throughput dual-colour VSVΔGpp screening system to discover novel inhibitors of specific viral glycoproteins. By simultaneously monitoring Zaire Ebola virus (EBOV) GP-driven infection and control infection mediated by VSV-G in the same assay, this approach enables rapid discrimination between selective inhibitors of Ebola virus entry and compounds causing non-specific antiviral activity, cytotoxicity, or general interference with VSV replication. Using this platform to screen a CytoSorb-derived human plasma compound library^17^, we identified Laudanosine, a degradation product of the clinically used neuromuscular blocker Atracurium, as a specific inhibitor of viral pathogens that depend on TPC2 endolysosomal trafficking for infection. Our findings demonstrate the power of human-derived compound libraries combined with viral entry screening approaches and identify Laudanosine as a host-directed antiviral against highly pathogenic late endosome-dependent viruses.

## RESULTS

To identify modulators of viral entry, we extracted crude filtrate from 6.5 kg of pooled CytoSorb (CSX) filters and separated it by chromatographic means (see methods). The extracted material consists of a complex mixture of hydrophobic, low to mid-molecular-weight plasma constituents (≤55 kDa), including inflammatory mediators, microbial components, endogenous metabolites, and adsorbed small-molecule drugs^18^. VSVΔG(EBOV-GP)pp expressing green fluorescent protein (GFP) and VSVΔG(VSV-G)pp expressing blue fluorescent protein (BFP) were titrated against each other and mixed to achieve a multiplicity of infection (MOI) of 0.1 for each pp on the target cells. The corresponding library fractions were added to human embryonic kidney (HEK293T) cells, which were subsequently inoculated with an equal dose of pseudovirus (Fig. 1a and Supplementary Fig. 1a). CSX fractions 17/18 (F17/18) restricted EBOV-GP-mediated entry by up to 80% while having no effect on VSV-G-mediated infection (Fig. 1b). To purify the inhibitor of EBOV-GP-dependent entry, CSX F17/18 were pooled and further separated chromatographically in an iterative process. Several fractions generated by this further separation specifically inhibited EBOV-GP-dependent cell entry (Fig. 1c; Supplementary Fig. 1b). Fraction 22, following fraction 16 were chosen for further purification because they were highly active and selective. The active subfraction CSX F17/18 F22, F16, F31 displayed a distinct peak in the A260/280 chromatographic spectrum (Fig. 1c and Supplementary Fig. 1b). MALDI-TOF-MS analysis revealed the decay pattern of Laudanosine, a 357.44 g/mol autocatalytically formed degradation product of the FDA-approved muscle relaxant Atracurium^19^, as the predominant molecule in this active fraction (Fig. 1d). To validate its antiviral activity, we tested commercial Laudanosine and found that it dose-dependently inhibits VSVΔG(EBOV-GP)-mediated entry into HEK293T cells, with an IC₅₀ of 27 µg/ml, while VSVΔG(VSV-G) infection remained unaffected (Fig. 1e). Fluorescence microscopy confirmed that Laudanosine selectively reduced EBOV-GP-mediated infection.

**Figure 1:**
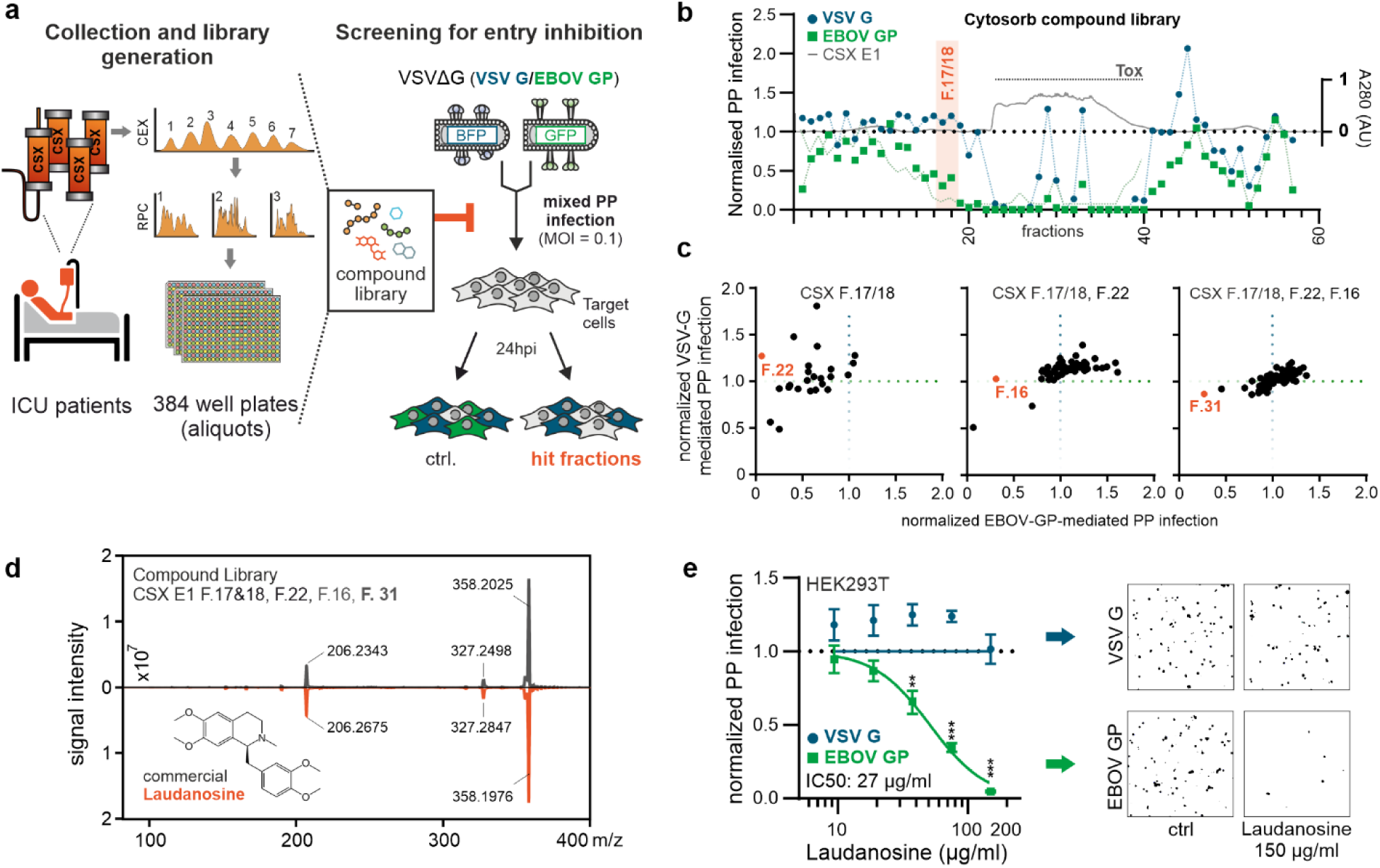
Identification of Laudanosine as an inhibitor of EBOV-GP-mediated entry. **a)** Workflow for generation of a compound library from patient samples and screening strategy. Fractions were generated by cation-exchange (CEX) and reverse-phase chromatography (RPC), aliquoted into 384-well plates, and tested for inhibition of viral cell entry using a VSVΔG pseudoparticle system bearing either EBOV-GP or VSV-G, with GFP/BFP reporters. **b)** Primary screen of Cytosorb Library showing normalized infection for VSVΔG bearing either EBOV-GP (green) and or VSV-G (blue). Fractions F. 17 and F. 18 specifically inhibit EBOV-GP-mediated infection with minimal effect on VSV-G; A280 trace indicates protein content. The initial screen was performed in biological singlets. **c)** Scatter plots of EBOV-GP-mediated versus VSV-G mediated cell entry for individual fractions. From left to right: Iterative subfractionation of CSX F17/18 F22, F16, F31, active fractions (red) were selected as hit and subjected to MS. Data represents the means ± SEM from n=3 **d)** Mass spectrometry profile of CSX F17/18, F22, F16, F31 (upper profile, black) leading to identification of Laudanosine in comparison to the profile of commercially available Laudanosine (lower profile, red). **e)** Dose-response analysis of Laudanosine on viral cell entry. Binaries of VSVΔG pseudotyped with VSV-G or EBOV-GP in the presence of DMSO control or 150 μg/mL Laudanosine. *, p < 0.05; **, p < 0.01; ***, p < 0.001.

EBOV entry depends on acidification-driven fusion during late endolysosomal trafficking^20,21^. Next, we tested whether Laudanosine also inhibits entry mediated by the GPs of other late endosome-dependent viruses in hepatocytes, the main target cells for EBOV replication (Fig. 2a). Entry mediated by EBOV, MARV (Marburg virus), LCMV and LASV GPs was inhibited with IC_50_ values of 23.9, 32.2, 61.4 and 73.6 µg/ml, respectively. In contrast, VSV-G-mediated entry that occurs mainly in early endosomes was only marginally reduced at the highest dose of 150 μg/ml of Laudanosine.

**Figure 2:**
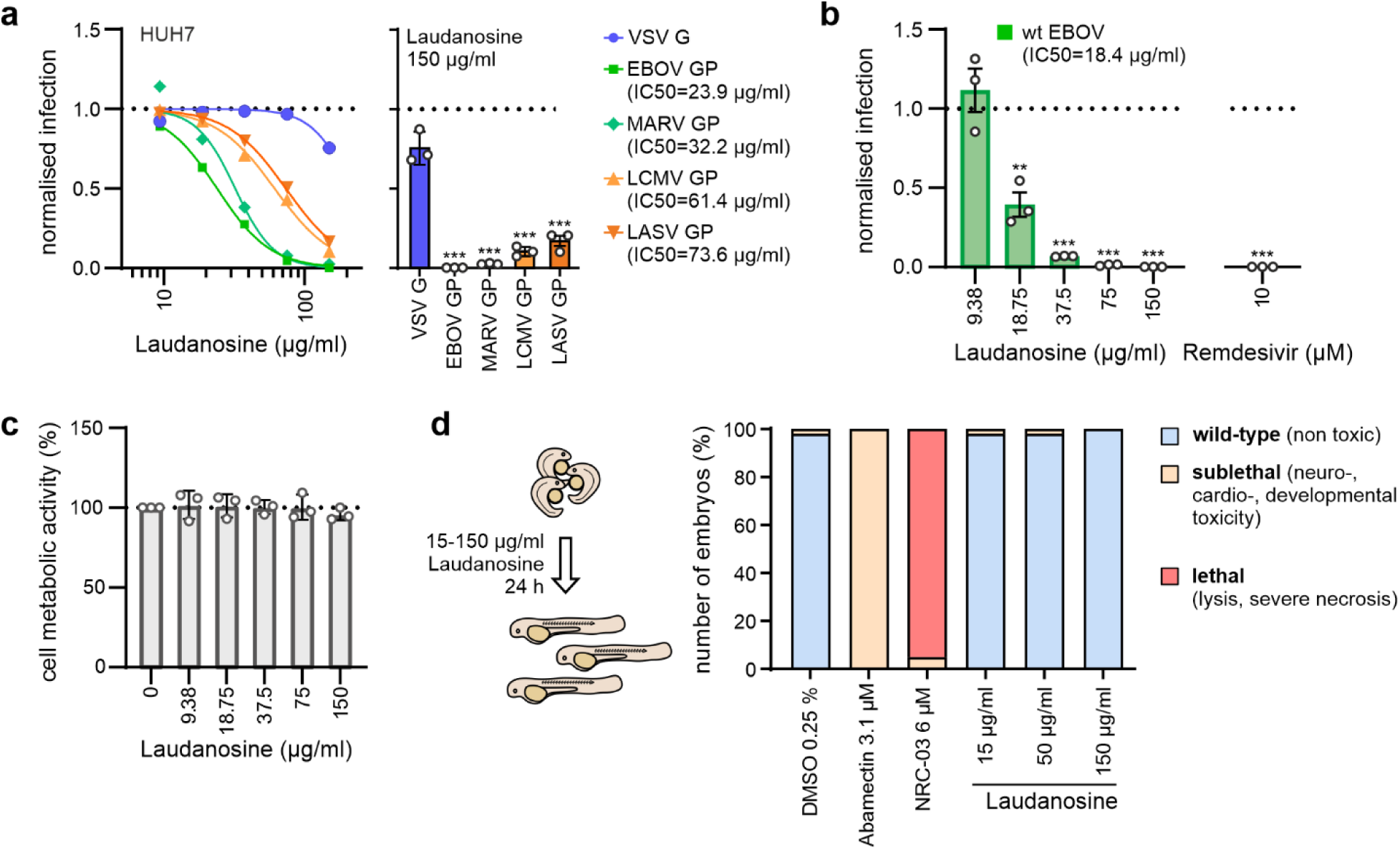
Effects of laudanosine on viral glycoprotein-mediated entry, EBOV infection, and cell viability. **a)** Dose-response analysis of Laudanosine-mediated inhibition of VSVΔG pseudoparticles bearing GPs from the indicated enveloped viruses in HuH7 cells. Data represents the means ± SEM from n=3 **b)** Antiviral activity of Laudanosine against authentic EBOV. HuH7 cells were infected with EBOV in the presence of increasing concentrations of Laudanosine, and viral RNA levels in infected cells were quantified by qRT-PCR 48 h post-infection. Data are normalised to DMSO-treated infected controls. Data represents the means ± SEM from n=3 **c)** Cell viability/metabolic activity of HuH7 cells treated with increasing concentrations of Laudanosine, showing no major cytotoxic effects at concentrations that inhibit viral entry. Data reptesents the means ± SEM from n=3 **d)** Laudanosine is not toxic to embryonic zebrafish. Twenty-four hours post fertilization, dechorionated zebrafish embryos were exposed to DMSO, Abamectin (neurotoxic control), NRC-03 (cytotoxic control) or increasing amounts of Laudanosine for 24 h. Data shown are derived from 60 embryos per group, sampled in two independent experiments. *, p < 0.05; **, p < 0.01; ***, p < 0.001

VSVΔG inhibition assays are a well-established surrogate system to study the entry of highly pathogenic viruses in low BSL environments^22–24^. However, they may not fully reproduce the mechanisms of wild-type virus infection (such as membrane composition, shape, or protein incorporation during budding)^25^. Thus, we determined the endpoint concentration of viral RNA in wild-type EBOV-infected cells. Laudanosine inhibited genuine EBOV infection with an IC₅₀ of 18.36 μg/mL (51.7 µM) (Fig. 2b)^26^, without showing cytotoxic effects (Fig. 2c). To assess potential toxic effects of Laudanosine *in vivo*, we used zebrafish embryos. This model is increasingly used in toxicity studies^27^. The transparency of the embryos allows not only evaluation of mortality, but also sublethal cytotoxicity (necrosis, lysis), developmental toxicity (developmental delay, malformations), cardiotoxicity (pericardial oedema, reduced circulation), and neurotoxicity (impaired touch escape response)^28^ (Fig. 2d). Laudanosine exposure did not display significant toxic effects in the zebrafish model, whereas NRC-03 (cytotoxic control) and Abamectin (neurotoxic control) exhibited the expected toxicities. Our finding that Laudanosine was nontoxic at concentrations 10-fold higher than the IC_50_ against wild-type EBOV suggests that it might be well tolerated *in vivo*.

Filoviruses depend on TPC2 for endosomal trafficking and fusion, thereby enabling release of viral nucleocapsids into the cytoplasm^29^. To assess the underlying antiviral mechanism, we analyzed two inhibitors of this pathway, Tetrandrine and SG-094, a simplified synthetic Tetrandrine analogue developed to retain TPC2 inhibition^30,31^. Both inhibited infection mediated by EBOV, MARV, LASV or LCMV GPs in VSVΔGpp entry assays (Supplementary Fig. 2a, 2b and Supplementary Table 1). To assess whether Laudanosine may interfere with TPC2 in a manner similar to Tetrandrin and SG-094, we performed molecular dynamics simulations using GROMACS 2022.2 with the CHARMM36 force field and the SG094-bound TPC2 structure as a reference model (PDB: 8OUO). Upon placement of Laudanosine in proximity to the allosteric binding pocket, the ligand underwent an initial reorientation and movement, reflected by a higher ligand Root Mean Square Deviation (RMSD) relative to the starting pose. Following this adjustment, the RMSD stabilized, indicating a sustained binding conformation. The average ligand RMSD, calculated after alignment to the protein backbone using the first frame as reference, was

0.575 ± 0.023 nm over the final 8 ns of the simulation. In comparison, SG-094 exhibited a lower average RMSD of 0.327 ± 0.047 nm, consistent with a preestablished binding pose closer to the ideal conformation. Apart from the expected structural rearrangement observed for Laudanosine, the low fluctuation after equilibration indicates a stable binding conformation. No dissociation events were observed for either ligand, supporting stable association with the binding site over the simulated timescale (Fig. 3b). To compare ligand–protein interactions, time-averaged short-range Lennard-Jones (LJ) and Coulomb interaction energies were computed for residues within 0.45 nm of each ligand over the final 8 ns of the simulation. Block averaging using 1 ns segments was applied to assess the robustness of the calculated interaction energies. The total ligand-protein interaction energies for SG-094 and Laudanosine were −196.0 ± 11.2 kJ/mol and −134.1 ± 11.8 kJ/mol, respectively. Autocorrelation analysis indicated effective sample sizes of approximately five to six independent blocks for both systems (Fig. 3b,c). Overall, these simulations suggest that Laudanosine stably associates with the allosteric inhibitory pocket in the VSD II region of TPC2.

**Figure 3:**
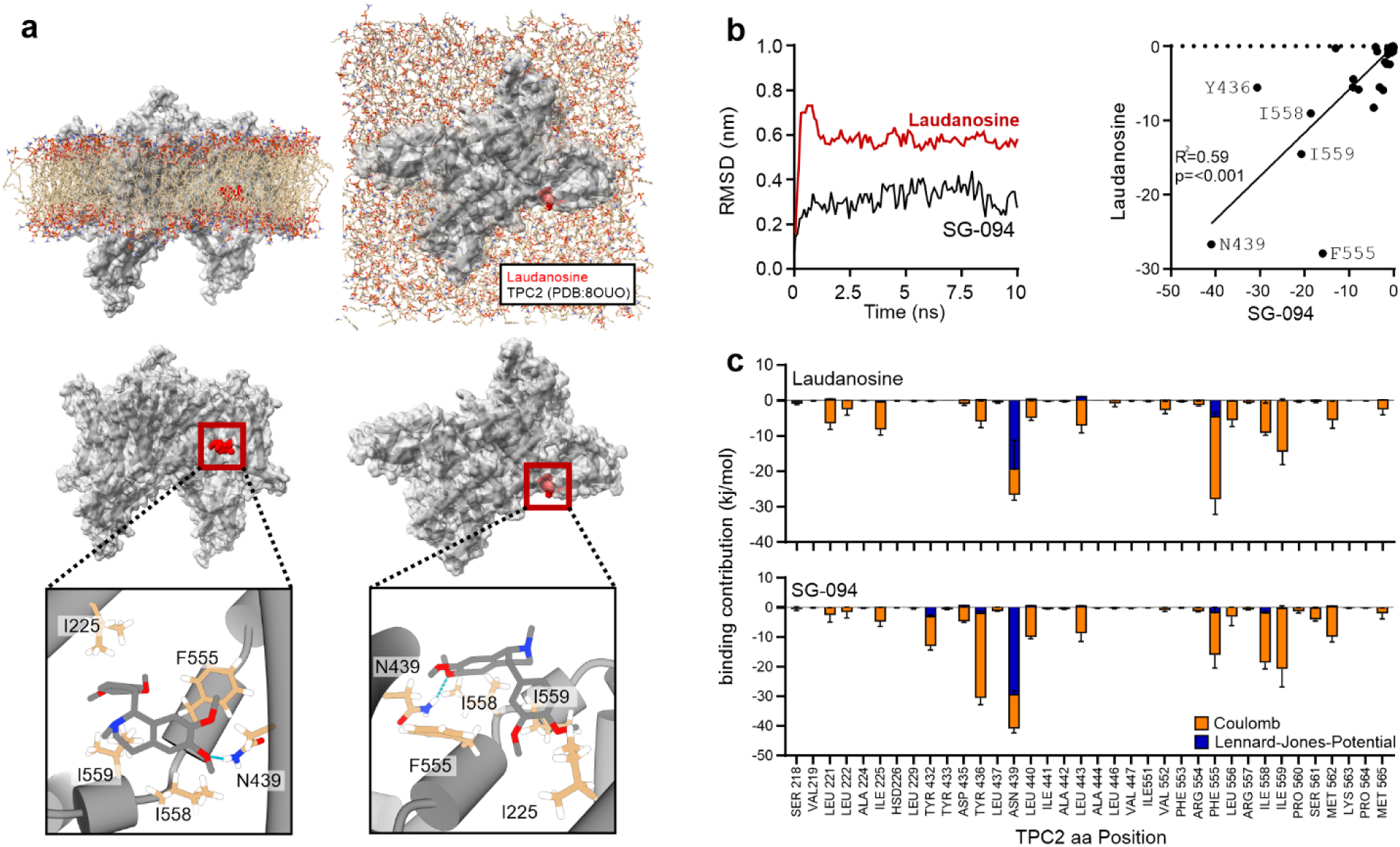
Predicted interaction of laudanosine with the TPC2 allosteric inhibitory pocket. **a)** Structural representation of TPC2 showing the allosteric inhibitory pocket within the voltage-sensor domain II region. SG-094 was used as reference ligand and Laudanosine was positioned into the same experimentally defined binding pocket. **b)** Ligand RMSD analysis of SG094 and Laudanosine during molecular dynamics simulations. Laudanosine initially reorients within the binding pocket, resulting in a higher RMSD relative to the starting pose, but subsequently stabilises over the simulated timescale. SG-094 remains closer to its initial binding pose. No dissociation events were observed for either ligand. **c)** Time-averaged short-range Lennard-Jones and Coulomb ligand-protein interaction energies calculated for residues within 0.45 nm of each ligand over the final 8 ns of the simulation. Total interaction energies indicate stable association of both ligands with the TPC2 binding pocket, with stronger interaction energy observed for SG094 than for Laudanosine. Data are shown as mean ± SD from 1 ns block averaging over the final 8 ns of the simulation. Molecular dynamics simulations were performed using GROMACS 2022.2 with the CHARMM36 force field and the SG094-bound TPC2 structure as reference model.

Occupation of the VSD II region of TPC2 has been associated with the functional impairment of ion transport leading to perturbed endosome-lysosome fusion and reduced autophagic flux, thereby causing accumulation of autophagosomes in the cytoplasm. To assess potential TPC2 blockage, we determined lysosome accumulation in HEK293T cells (Fig. 4a) and utilized HEK293T autophagy reporter cells, stably expressing GFP labelled Microtubule-associated proteins 1A/1B light chain 3B (GFP-LC3B). Membrane-associated LC3B as a marker of autophagosomes, was revealed by mild permeabilization of the cells and washing out soluble LC3B^32^. Treatment with increasing concentrations of Laudanosine resulted in a dose-dependent increase in membrane-associated GFP-LC3B signal, indicating increased autophagosome numbers (Fig. 4b). Co-treatment with saturating concentrations of Bafilomycin A1, which blocks autophagosome-lysosome fusion and lysosomal acidification, did not further increase GFP-LC3B accumulation in Laudanosine-treated cells (Fig. 4c, d). This suggests that Laudanosine and Bafilomycin A1 interfere with a late step of autophagic flux, like Bafilomycin A1. Increased processing of LC3B into lipidated LC3B-II and elevated levels of the autophagy receptor p62 are further hallmarks of impaired autophagic turnover. Immunoblot analysis revealed accumulation of LC3B-II and p62/SQSTM1 upon Laudanosine treatment (Fig. 4d). Together, these data indicate that Laudanosine impairs endolysosomal trafficking and autophagic flux, in line with inhibition of TPC2-dependent ion conductance.

**Figure 4:**
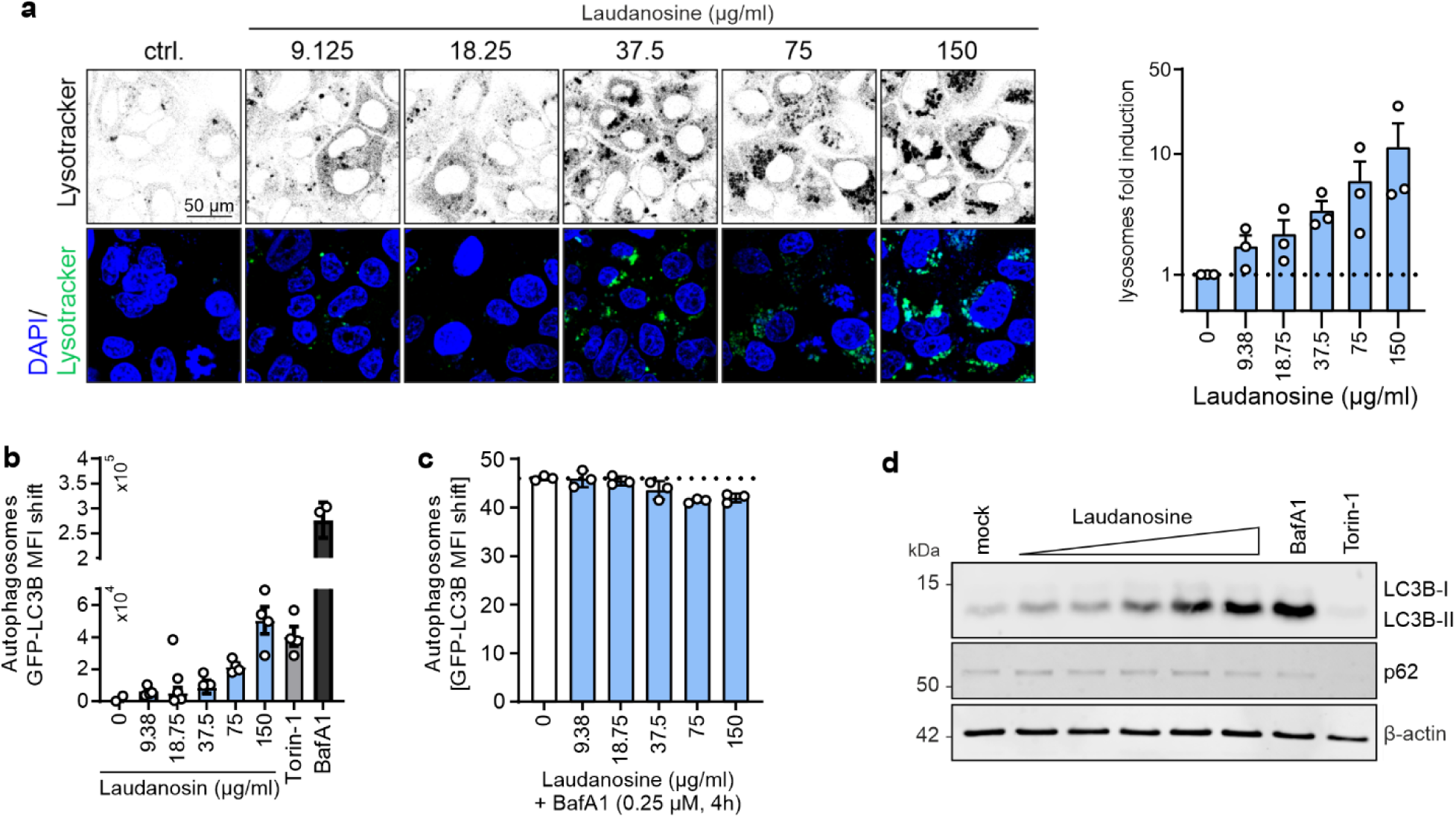
Laudanosine impairs autophagic flux. **a)** Representative confocal images of Huh7 cells treated for 24 h with increasing concentrations of Laudanosine (9.125-150 µg/ml) or DMSO control and stained with LysoTracker (green) and DAPI (blue). Images were acquired under identical settings. Laudanosine induced a dose-dependent accumulation of lysosomes. Right: quantification of lysosomal increase relative to DMSO control. Data represents the means ± SEM from n=3 **b)** Quantification of membrane-associated GFP-LC3B in HEK293T-GL autophagy reporter cells treated with increasing concentrations of Laudanosine. Cells were permeabilised to remove soluble GFP-LC3B, and membrane-bound GFP-LC3B was quantified by flow cytometry as a measure of autophagosome accumulation. Data represents the means ± SEM from n=4 **c)** Autophagic flux analysis in HEK293T-GL cells treated with increasing concentrations of Laudanosine in the presence or absence of bafilomycin A1 (BafA1). Co-treatment with BafA1 did not further increase GFP-LC3B accumulation in Laudanosine-treated cells. Data represents the means ± SEM from n=3 **d**) Representative immunoblot analysis of autophagy markers in Laudanosine-treated cells. Accumulation of LC3-II and p62/SQSTM1 is consistent with reduced autophagosomal degradation. *, *p* < 0.05; **, *p* < 0.01; ***, *p* < 0.001.

## DISCUSSION

Filoviruses remain major global health threats because they combine high pathogenicity, recurrent outbreaks and limited therapeutic options. Although Ebola virus disease occurs predominantly in sporadic outbreaks, these events are frequently associated with severe disease and high case fatality rates^33,34^. The devastating 2013-2016 West African Ebola epidemic caused by Zaire ebolavirus (Makona variant), resulting in ∼28,600 infections and 11,300 deaths^35^, together with the ongoing outbreak caused by Bundibugyo virus in the Democratic Republic of the Congo and Uganda, with more than 330 confirmed cases to date (June 2026), underscore the urgent need for antivirals against ebolaviruses and related pathogens^36,37^. Here, we identify Laudanosine as an inhibitor of TPC2-dependent endolysosomal trafficking that restricts viral entry through late endosomal pathways. Laudanosine selectively inhibited entry mediated by Ebola, Marburg, LCMV and Lassa virus envelope glycoproteins, suppressed authentic EBOV infection without detectable toxicity in cultured cells or embryonic zebrafish, and recapitulated key features of impaired endolysosomal function.

The discovery of Laudanosine emerged from a screening strategy designed to exploit human-derived libraries for antiviral agents^3,18^. CytoSorb-derived libraries extend this concept by representing complex mixtures of circulating peptides, metabolites, inflammatory mediators, administered drugs and their derivatives from severely ill patients^18,38,39^. To isolate selective modulators of viral entry from this heterogeneous source, we established a high-throughput dual-colour VSVΔGpp screening platform. This approach allowed us to simultaneously monitor EBOV-GP-mediated entry and VSV-G-dependent infection within the same well and to discriminate between selective inhibitors or compounds causing cytotoxicity or broad non-specific effects. Since VSV-G mediates fusion predominantly in early endosomes, whereas filovirus and arenavirus glycoproteins require late endosomal maturation^40^, the assay allows to uncover host-directed entry inhibitors. In addition, this platform is scalable and could be expanded by incorporating additional fluorescent reporters and viral glycoproteins to identify antiviral compounds targeting diverse viral pathogens.

Laudanosine is a degradation product of Atracurium, a benzylisoquinolinium neuromuscular blocking agent widely used to induce muscle relaxation during anaesthesia and intensive care treatment^41–43^. Its identification in CytoSorb-derived fractions is consistent with the clinical application of haemoadsorption devices in critically ill patients suffering from severe systemic inflammation, sepsis, acute respiratory distress syndrome or severe viral infections^44^. Many of these patients require prolonged mechanical ventilation and frequently receive neuromuscular blocking agents such as Atracurium to improve ventilator synchrony and facilitate lung-protective ventilation. Consequently, metabolites like Laudanosine may accumulate among the plasma-derived compounds captured by CytoSorb cartridges. Previous studies in mechanically ventilated patients receiving continuous Atracurium infusion over 72 hours (0.5 mg/kg/hour) reported plasma Laudanosine concentrations of up to 20 µg/ml without specific adverse effects attributable to this metabolite^43^. Notably, the IC₅₀ observed against authentic EBOV infection (18.36 µg/ml) falls within this concentration range^43,45^. Although additional pharmacokinetic analyses and *in vivo* efficacy studies are required, our findings support the potential use of laudanosine as a therapeutic agent to treat acute Ebola virus infection.

Mechanistically, our results support that Laudanosine restricts viral entry by interfering with TPC2-dependent endolysosomal trafficking. TPC2 regulates ion fluxes required for endosome maturation and fusion events exploited by filo- and arenaviruses. Molecular dynamics simulations predicted stable association of Laudanosine with the allosteric inhibitory pocket of TPC2^31,46,47^. Consistent with impaired TPC2 function, Laudanosine induced accumulation of lysosomal compartments and GFP-LC3B-positive autophagosomal structures. Co-treatment with Bafilomycin A1 did not further enhance this phenotype, indicating that Laudanosine interferes with late stages of autophagic flux rather than increasing autophagosome formation^48–51^. Accumulation of LC3-II and p62/SQSTM1 confirmed impaired autophagosomal turnover. Thus, Laudanosine mirrors key aspects of defective autophagosome–lysosome and endosome-lysosome fusion^51,52^, revealing an unexpected biological activity beyond its known role as an Atracurium metabolite^31,46,47^.

In summary, the combination of human-derived compound screening, dual-colour VSVΔGpp entry assays, authentic EBOV validation, molecular modelling and functional autophagy analyses identifies Laudanosine as a previously unrecognised modulator of TPC2-dependent endolysosomal trafficking. These findings highlight late endosomal trafficking as a druggable vulnerability shared by several highly pathogenic viruses and demonstrate the potential of mining human-derived compound libraries for antiviral discovery. Further analyses of Laudanosine or derivatives thereof may lead to the development of host-directed antivirals against filoviruses and other viral pathogens dependent on endolysosomal entry pathways.

## METHODS

### Cell culture and viruses

All cells were cultured at 37°C in a 5% CO2 atmosphere with 90% humidity. Human embryonic kidney 293T cells purchased from American type culture collection (ATCC: #CRL3216), autophagy reporter HEK293T cells stably expressing GFP-LC3B (GL)^32^ cells and Human hepatoma-derived HuH-7 cells were cultivated in Dulbecco’s Modified Eagle Medium (DMEM, Gibco) supplemented with 10% (v/v) heat-inactivated fetal bovine serum (FBS, Capricorn), 2 mM L-glutamine (PANBiotech), 100 µg/ml streptomycin (PANBiotech) and 100 U/ml penicillin (PANBiotech). I1-Hybridoma cells were purchased from ATCC (#CRL-2700) and cultured in Roswell Park Memorial Institute (RPMI) medium supplemented with 10% (v/v) heat-inactivated FBS (Capricorn), 2 mM L-glutamine (PANBiotech), 100 µg/ml streptomycin (PANBiotech) and 100 U/ml penicillin (PANBiotech).

Pseudoparticle production. HEK 293T cells (1 × 10⁶ per well) were seeded into 6-well plates (Sarstedt) 18 h prior to transfection. Cells were transfected with 5 µg of a GP expression plasmid using the calcium phosphate method. At 48 h post-transfection, cells were infected with VSVΔG-GFP or VSVΔG-BFP particles pseudotyped with VSV-G at a MOI of 3. 16 h post-infection, pseudotyped VSVΔG-GFP/BFP particles were harvested from the culture supernatant. Cells were removed by two consecutive centrifugation steps at 4,000 × g for 5 min at 20 °C. For pseudotypes bearing GPs other than VSV-G, residual input particles carrying VSV-G were neutralized by adding 10% (v/v) I1 hybridoma supernatant (I1, mouse hybridoma supernatant, CRL-2700; ATCC) to the harvested supernatant. Pseudotyped particles were subsequently stored at −80 °C until further use.

### Library generation and sample collection

Twenty CytoSorb (CSX) cartridges obtained from different donors were collected at Ulm University Medical Center under approval of the ethics committee vote 150/16. Bound material was extracted with acetonitrile, and the pooled extract was diluted with demineralized water to a final concentration of 5% acetonitrile. The extract was acidified with HCl to pH 2.6 and loaded onto a reverse-phase column (Sepax PolyRP). Compounds were eluted using a gradient from 5% to 100% acetonitrile. In total, 55 fractions were generated and subsequently tested for antiviral activity. The hemofiltrate library was generated in a previous work described^53^. The Ethic approval for the generation of peptide libraries from hemofiltrate was obtained from the Ethics Committee of Ulm University (91/17).

### Infection with Zaire Ebola virus

Experiments involving EBOV were conducted at the BSL-4 facility at the Philipps University Marburg in accordance with national and international regulations. EBOV (GenBank accession number NC_002549) was propagated as virus stock in VeroE6 cells. Viral titers of stock virus were calculated based on plaque-forming units (PFU) in VeroE6 cells. HuH7 cells (1x 104 cells/ 96well) were infected with EBOV in serum-free medium using a multiplicity of infection (MOI) of 0.1. The virus- containing supernatant was discharged 1 hour post infection (hpi) and replaced with medium (containing 3% FCS, penicillin (50 units/ml) and streptomycin (50 mg/ml)) and supplemented with corresponding Laudanosin as indicated. Remdesivir (10µM in DMSO) was included as further control. Cells were incubated for 48 hours at 37 °C and 5% CO2.

### RNA Isolation and qRT PCR

To extract RNA from cell lysate, remaining supernatant was removed from the infected cells and 130 µl RLT-buffer (RNeasy Mini Kit, Qiagen) with 1.3 µl 2-Mercaptoethanol and 0.5 µl Bromphenol blue were added. The plate was incubated for 10 minutes, followed by the addition of 130 µl 70% ethanol. Inactivated samples were transferred to a new 96-well plate and sealed with an aluminum foil, disinfected according to BSL-4 regulations and subsequently transferred to BSL-1 lab for further processing.

RNA was purified using a fully automatic nucleic acid extraction system (Tecan). Briefly, 200 µl sample was combined with 200 µl of DNA/RNA Shield™ in a deep-well plate and purified through the utilization of the Quick-DNA/RNA viral MagBead kit (Zymo research). The kit was used according to the manufacturer’s instructions with the following adaptations: 20 µl magnetic beads, 460 µl of wash solutions and 460 µl ethanol were used and RNA was eluted from the magnetic beads by the addition of 50 µl DNase/RNase-Free Water. EBOV RNA isolated from cells was analysed using RealStar® Filovirus Screen RT-PCR Kit 1.0 (targeting L gene) [altona Diagnostics]. The kits were used according to the manufacturer’s instructions with the following adaptations: 2.5 µl Master A and 7.5 µL Master B solution were mixed with 5 µl sample from RNA extraction. A positive control RNA was incorporated into each plate. All RT-PCRs shown here were conducted for a total of 45 cycles.

### MALDI-TOF analysis

Samples were analyzed by an Axima Confidence MALDI-TOF MS (Shimadzu, Japan) in positive linear mode on a 384-spot stainless-steel sample plate. Spots were coated with 1 µL of 5 mg/mL CHCA dissolved in matrix diluent (Shimadzu, Japan), and the solvent was allowed to air-dry. Then, 0.5 µL of sample or standard was applied to the dry, pre-coated well and immediately mixed with 0.5 µL of matrix; the solvent was allowed to air dry. All spectra were acquired in the positive ion linear mode using a 337-nm N2 laser. Ions were accelerated from the source at 20 kV. A hundred profiles were acquired per sample, and 20 shots were accumulated per profile. The equipment was calibrated with a standard mixture provided in the TOFMixTM MALDI kit (Shimadzu, Japan). Measurements and MS data processing were controlled by the MALDI-MS Application Shimadzu Biotech Launchpad 2.9.8.1 (Shimadzu, Japan).

Structural elucidation by MS/MS. Fragmentation experiments were performed on the same instrument and under the acquisition conditions described above, with the following modifications. Precursor ions were fragmented by post-source decay (PSD), and the laser power was increased by 50% relative to the MS settings. The ion gate was activated with a precursor selection window of ±0.5 Da. A total of 1000 profiles were acquired per sample, with 20 laser shots accumulated per profile. Fragment ion spectra were searched against the MetFrag in silico fragmentation tool and the MassBank of North America (MoNA) experimental spectral library. Candidate identifications were then confirmed by acquiring the PSD spectrum of the corresponding commercial standard under identical conditions and comparing it to the sample spectrum. Spectral similarity between the sample and the commercial standard fragment spectra was quantified as cosine score using the CosineGreedy algorithm implemented in the matchms Python library (v0.32.0)^54^, with a fragment m/z tolerance of 0.1 Da, m/z power = 0 and intensity power = 1 (Similarity 0.9943). The analysis notebook is available as Supplementary Material at https://doi.org/10.5281/zenodo.19697427

### Dual-colour screen

To perform the dual-colour screen, 10 µL of each library fraction was transferred into 384-well plates (Corning). Subsequently, 30 µL of freshly generated VSVΔG(EBOV-GP)-GFP and VSVΔG(VSV-G)-BFP pseudotyped particles were added at a MOI of 0.1. HEK 293T cells (5 × 10⁴ per well) were then added to a final volume of 70 µL. Plates were incubated for 18 h at 37 °C in a humidified incubator (90% humidity, 5% CO₂). Following incubation, GFP/BFP-positive cells were automatically quantified using a Cytation 3 microplate reader (BioTek). Hit fractions were subsequently validated by retesting in triplicate using the same assay conditions. Confirmed active fractions were subjected to further subfractionation and analysis.

### Cytotoxicity Assay

Cell viability of Huh7 cells treated with Laudanosine (EDQM) or Tetrandrine (Targetmol) was determined using the CellTiter-Glo® kit (Promega) according to the manufacturer’s instructions. In short, 25.000 Huh7 cells per well were seeded in 96-well format and treated with different concentrations of Laudanosine. 24 h after treatment, 100 µl of CellTiter-Glo® 2.0 reagent was added to the cells and luminescence was quantified using a Cytation 3 microplate reader (BioTek).

### Laser Scanning Microscopy

To monitor the effects of Laudanosine on lysosomal compartments, Huh7 cells were seeded in 8-well µ-Slides (ibidi) with an ibiTreat surface. Cells were incubated for 24 h with increasing concentrations of Laudanosine or DMSO as a vehicle control. Subsequently, cells were stained by addition of LysoTracker Green DND-26 (ThermoFisher, 1:100 dilution) and DAPI (Manufacturer), followed by incubation for 30 min at 37 °C (5% CO₂, humidified atmosphere). Cells were washed three times with culture medium prior to image acquisition. Images were acquired using an inverted confocal microscope (Zeiss 710) Fluorescence signals for DAPI (blue channel) and LysoTracker (green channel) were recorded sequentially using constant excitation and emission settings across all samples. For quantitative analysis, randomly selected fields of view were imaged and z-stacks were acquired under identical settings.

### Zebrafish

For in vivo studies, wild-type zebrafish embryos (Danio rerio) were dechorionated at 24 h post fertilization (hpf) using digestion with 1 mg/mL Pronase (Sigma) in E3 medium (83 μM NaCl, 2.8 μM KCl, 5.5 μM 202 CaCl_2_, 5.5 μM MgSO_4_). In a 96-well plate, 3 embryos per well were exposed for 24 h to 100 μL of E3 containing Laudanosine at the concentrations indicated in the figures. Two independent assays were performed, each with 10 × 3 embryos. The solvent (DMSO), diluted in E3, was used as negative control at the same amount as introduced by the Laudanosine stock. As positive control for acute toxicity/cytotoxicity the pleurocidin antimicrobial peptide NRC-03 (GRRKRKWLRRIGKGV-KIIGG AALDHL-NH2) was used at a concentration of 6 μM as described^55^. Abamectin at a concentration of 3.125 μM was used as positive control for neurotoxicity^56^. At 48 hpf (after 24 h of incubation) embryos were scored in a stereomicroscope for signs of cytotoxicity (lysis and/or necrosis), developmental toxicity (delay and/or malformations) or cardiotoxicity (heart oedema and/or reduced or absent circulation). Each embryo was also touched with a needle and reduced or absent touch response (escape movements) was evaluated as signs of neurotoxicity if and only if no signs of acute toxicity were present in the same embryo. Embryos were categorized within each of these toxicity categories into several classes of severity. Fisher’s exact test was used to calculate whether the distribution of embryos into toxicity classes differed significantly between the PBS negative control and the test substances.

### Autophagosome measurement by flow cytometry

Autophagy reporter cells were analyzed by flow cytometry as previously described^32,57^. In brief, HEK293T cells stably expressing GFP-LC3B (GL) cells were seeded (5 x 104 per well of a 96-well F-bottom plate). The next day, the medium was removed and cells were stimulated with increasing amounts of Laudanosine (150, 75, 37.5, 18.75, 9.375 µg/ml), 1 µM Torin-1 or left untreated. Cells were optionally co-stimulated with 0.25 µM Bafilomycin A1. After 4 h, the supernatant was removed and the cells were detached using Trypsin/EDTA 0.05%/ 0.02%. The harvested cells were washed with DPBS and treated with DPBS containing 0.05% Saponin at 4°C for 20 min for permeabilization. After washing the cells twice with DPBS to wash out the non-membrane bound GFP-LC3B, cells were fixed in 2% paraformaldehyde (PFA). The mean fluorescence intensity (MFI) of membrane-bound GFP-LC3B was then measured by flow cytometry on a Cytek Northern Lights using the software Spectro and analyzed by FlowLogic. The MFI value of the control was used as baseline and subtracted.

### Whole-cell lysates

For preparing whole-cell lysates (WCL) cells were washed with DPBS, centrifuged for 4 min at 300 x g and lysed in NP40 lysis buffer (1% (v/v) IGEPAL CA-630, 2.5 M NaCl, 1 M HEPES pH 7.4, deionized water) supplemented with 1:500 protease inhibitor for 10 min on ice. Cell debris were pelleted by centrifugation for 20 min at 4 °C and 20,000 x g and the WCL was transferred to fresh tubes. Subsequently, the WCL was mixed with 6x protein sample loading buffer (187.5 mM Tris-HCl adjusted to pH 6.8, 75% (v/v) glycerol, 6 % (w/v) SDS, 0.3 % (w/v) Orange G) to a final concentration of 1x containing 10% (v/v) dithiothreitol (DTT) dissolved in deionized water and stored at -20 °C until use.

### SDS-PAGE and immunoblotting

For protein separation, whole-cell lysates were heated up to 95 °C for 10 min before use and loaded on 12% Bis-Tris gels. The gels were running in 1x MES-SDS running buffer (20x MES-SDS running buffer diluted in deionized water) 30 min at 30 V and subsequently for 80 min at 90 V. Next, the separated proteins were wet blotted onto an Immun-Blot PVDF Membrane using the Thermo Fisher Mini Blot Module system (Cat# B1000) according to the manufacturers instruction and using the transfer buffer (25 mM Tris and 192 mM glycine at a pH of 8.3 with 20% (v/v) methanol) at a constant voltage of 20 V for 1 h. After blocking in Blocker Casein in PBS for 1 h, the proteins on the membrane were stained with primary antibodies diluted in TBS-T (1x TBS with 0.05% (v/v) Tween 20 and 0.1% (v/v) Blocker Casein in PBS) for 2 h at RT or overnight at 4 °C. The following primary antibodies were used: 1:10,000 Monoclonal mouse anti-β-actin Antibody (AC-15), 1:1,000 Monoclonal rat anti-GAPDH Antibody (W17079A), 1:1,000 Monoclonal mouse anti-p62 Antibody (2C11), 1:200 Polyclonal rabbit anti-LC3 Antibody. After incubation with the primary antibodies, the membrane was washed three times with TBS-T for 5 min at RT. Subsequently, the membrane was incubated in IRDye secondary antibodies diluted in TBS-T for 30 min under shaking. The following IRDye secondary antibodies were used in this study: IRDye 800CW Goat anti-Mouse IgG, IRDye 680RD Goat anti-Rat IgG, Goat anti-rabbit IgG H&L (HRP). After three times washing with TBS-T, the fluorescent signal of the IRDye secondary antibodies or the HRP after incubating the membrane with Biorad Clarity Western ECL Substrate for 30 s was detected using a LI-COR Odyssey M (LI-COR, Lincoln, NE, US) and the Image Acquisition 2.2. Image processing and quantification of band intensities were analyzed by using the software Image Studio Version 6.

### Molecular dynamics simulation

Membrane-embedded TPC2-ligand systems were prepared using CHARMM-GUI and simulated with GROMACS using the supplied CHARMM force-field parameters and TIP3P water topology. The system containing Laudanosin comprised one copy of TPC2, the ligand itself, 395 DPPC lipids, 54,905 water molecules, 150 K+ ions, and 162 Cl- ions. The reference system comprised one copy each of TPC2, the original ligand SG_094, 395 DPPC lipids, 56,412 water molecules, 153 K+ ions, and 165 Cl- ions. Approximately 150 mM KCl was added, together with 12 additional chloride ions to neutralize the net positive charge of each system. The initial box dimensions were 13.01535 x 13.01535 x 14.95920 nm for the Laudanosin system and 13.01057 x 13.01057 x 15.16595 nm for the reference SG94 system.

Each system was energy-minimized for up to 5,000 steepest-descent steps using a convergence criterion of 1,000 kJ mol-1 nm-1. Where both minimizations converged before reaching the step limit. The systems were subsequently equilibrated for a total of 1.875 ns using the six-stage CHARMM-GUI membrane-equilibration protocol. The first three stages were performed for 125 ps each with a 1 fs time step, followed by three 500 ps stages with a 2 fs time step. Positional restraints on the protein backbone, protein side chains, and lipids, as well as protein dihedral restraints, were progressively reduced during equilibration. Temperature was maintained at 303.15 K using the velocity-rescaling thermostat with separate coupling groups for the solute, membrane, and solvent and a coupling constant of 1.0 ps. From the third equilibration stage onward, pressure was maintained at 1 bar using semi-isotropic C-rescale pressure coupling with a coupling constant of 5.0 ps and a compressibility of 4.5 ⋅ 10-5 bar-1.

Production simulations were performed for 10 ns as ten consecutive 1 ns segments using a 2 fs time step. Long-range electrostatic interactions were treated using particle-mesh Ewald summation with a real-space cutoff of 1.2 nm. Lennard-Jones interactions were force-switched between 1.0 and 1.2 nm. Bonds involving hydrogen atoms were constrained using LINCS. Coordinates were written every 100 ps, while energies and log-file quantities were recorded every 2 ps.

### Data Availability

The datasets generated during and/or analyzed during the current study are available from the corresponding authors on request. Source data are provided with this paper.

### Statistics

Statistical analyses were performed using GraphPad PRISM 10.4.1 (GraphPad Software). P-values were determined using a two-tailed unpaired Student’s t test. Unless otherwise stated, data are shown as the mean of at least three independent experiments ± SEM.

## Acknowledgments

We thank Kerstin Regensburger, Regina Burger, Jana-Romana Fischer, Birgit Ott, Martha Meyer, Nicole Schrott, Daniela Krnavek and Magdalena Weiß for technical assistance. T.S. and D.C. are part of the International Graduate school for Molecular Medicine (IGradU), Ulm. This study was supported by DFG grants to F.K. (CRC 1279, SPP 1923), K.M.J.S. (CRC1279, SP1600/6-1), G.W. (CRC1279) and T.J. (CRC 1279). Regarding the computational resources, support by the state of Baden-Württemberg through bwHPC and the DFG through grant no INST40/575-1 FUGG (JUSTUS-2 cluster) is gratefully acknowledged. F.Z. was funded by the “Bausteine” program of Ulm University (project number L.SBN.0225)

## Author Contributions

T.S. performed most experiments with the help of D.C. and L.T ; J.K., and C.J. T.J. performed molecular modelling analyses. F.K. and N.B. performed wild-type EBOV experiments. S.K. and K.M.J.S. performed experiments on autophagic flux. N.P., A.A.R.A., and L.S. performed compound purification. J.K. and G.W. performed zebrafish toxicity assays. F.Z. and F.K. conceived the study, planned experiments, and wrote the manuscript. All authors reviewed and approved the manuscript.

## Materials & Correspondence

Further information and requests for resources and reagents should be directed to and will be fulfilled by Fabian Zech.

