## Supplementary material for "Laudanosine restricts Ebola virus entry by targeting TPC2-dependent endolysosomal trafficking": Suplementary Figure 1

**Suplementary Figures**

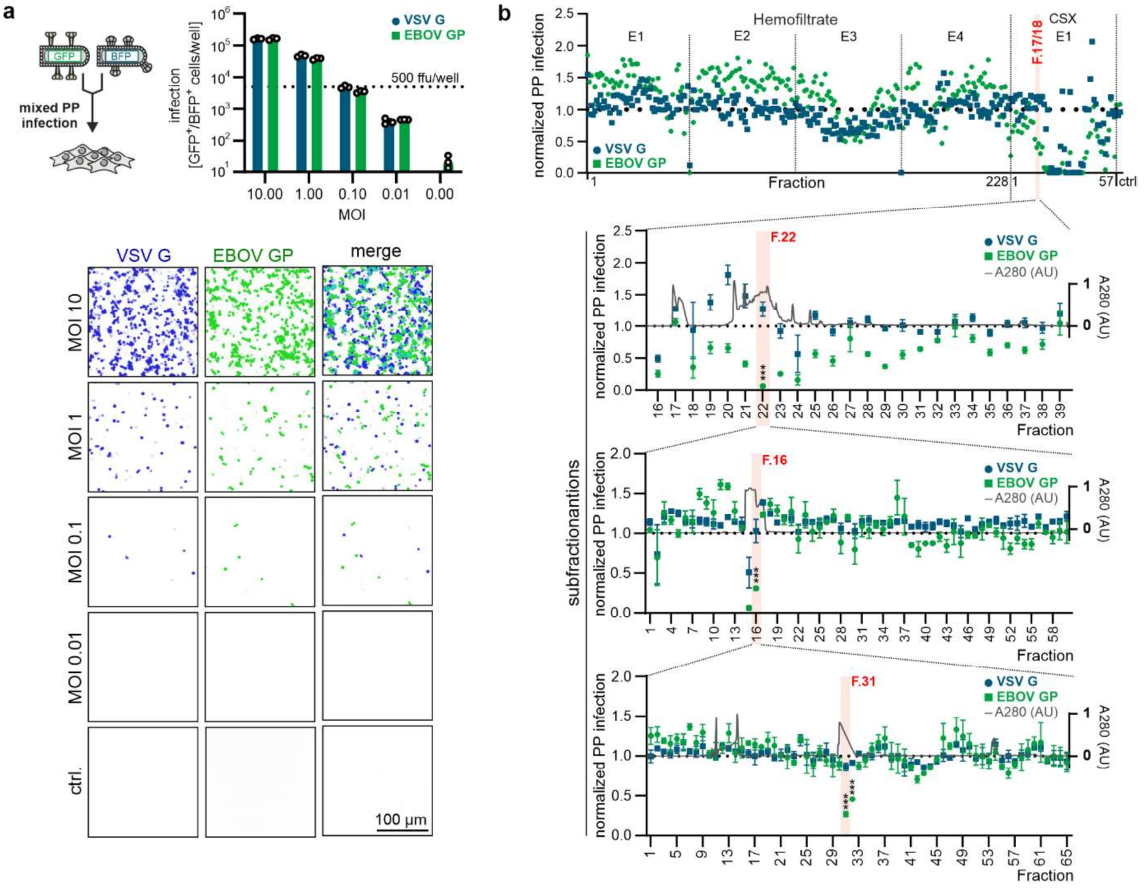


**Suplementary Figure 1: Dual-colour screening of human-derived compound libraries for modulators of EBOV-GP-mediated entry. a)** Dual-colour infection assay validation. HEK293T cells were co-infected with VSVΔG pseudoparticles encoding GFP or BFP and bearing EBOV-GP or VSV-G, respectively, across a range of MOIs (10-0.01). Infection levels correlate across both pseudotypes. Representative binary images of the titration series are shown below. Data represent the means ± SEM from n=3. **b)** Dual-colour screening workflow. Human-derived hemofiltrate fractions (E1-E4) and Cytosorb (CSX) fractions were screened in a dual-colour assay using HEK293T cells co-infected with EBOV-GP (green) and VSV-G (blue) VSVΔGpp in the same well. Most fractions affected infection rates similarly, with selective inhibition of EBOV-GP-mediated entry observed for CSX fraction F. 17/18 and retained through subfractions F. 22 and F. 16. Black trace indicates A280 (peptide abundance). The initial screen was performed in biological singlets, all sub fractionations represent the means ± SEM from n=3. *, $p < 0.05$; **, $p < 0.01$; ***, $p < 0.001$
