## Supplementary material for "Laudanosine restricts Ebola virus entry by targeting TPC2-dependent endolysosomal trafficking": Suplementary Figure 2

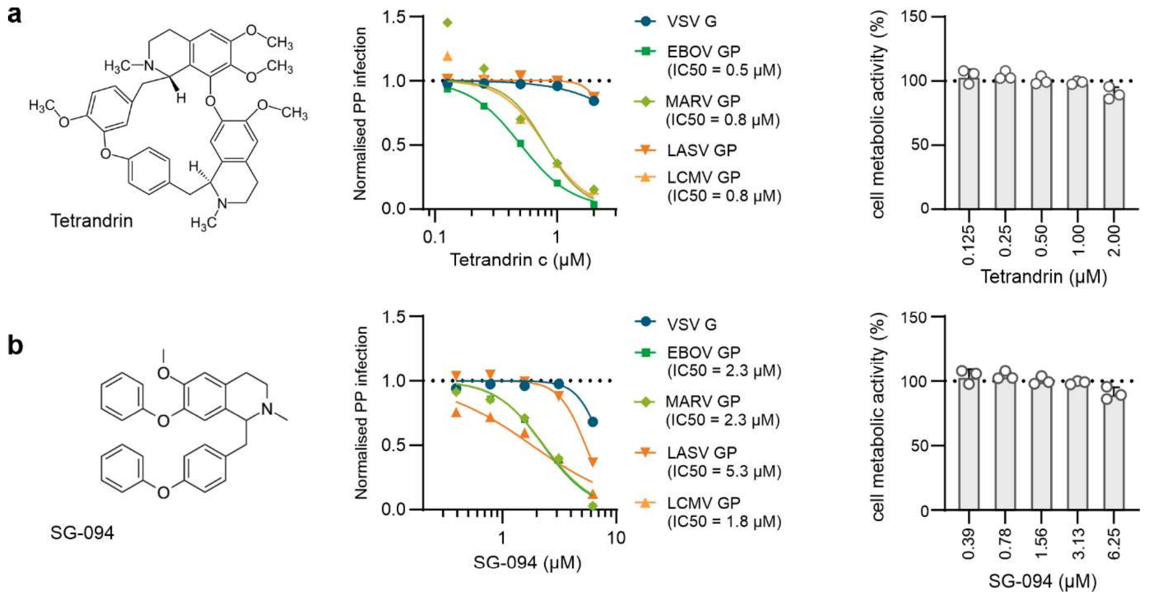


**Suplementary Figure 2: Effects of Tetrandrin and SG-094 on viral glycoprotein-mediated entry and cell viability. a)** Dose-response analysis of tetrandrin (upper part) and SG-094-mediated (lower part) inhibition of VSVΔG pseudoparticles bearing GPs from the indicated enveloped viruses in HuH7 cells. $IC_{50}$ values are indicated. Data represent the means ± SEM from n=3. **b)** Cell viability/metabolic activity of HuH7 cells treated with increasing concentrations of tetrandrin (upper part) or SG-094 (lower part). Data represent the means ± SEM from n=3.
