## Supplementary material for "Laudanosine restricts Ebola virus entry by targeting TPC2-dependent endolysosomal trafficking": Suplementary Table 1

**Suplementary Table 1: Inhibition of endolysosome dependent viral entry by TPC2 inhibitors**

| $IC_{50}$ | Laudanosin (µg/ml) | | Laudanosin (µM) | | Tetrandrin (µM) | SG-094 (µM) |
|---|---|---|---|---|---|---|
| | HEK293T | HUH 7 | HEK293T | HUH 7 | HUH 7 | HUH 7 |
| **EBOV GP** | 27.00 | 23.93 | 76.00 | 67.33 | 0.49 | 2.30 |
| **MARV GP** | | 32.19 | | 90.57 | 0.79 | 2.30 |
| **LCMV GP** | | 61.35 | | 172.61 | 0.79 | 5.30 |
| **LASV GP** | | 73.63 | | 207.16 | >2 | >5 |
| **VSV G** | >150 | 150.00 | >422 | >422 | >2 | >5 |
| **WT EBOV** | | 18.36 | | 51.66 | | |
